# Genomics reveals population structure despite high connectivity of common sole, *Solea solea*, and European plaice, *Pleuronectes platessa*, in the Celtic Sea and western English Channel

**DOI:** 10.64898/2026.06.10.731368

**Authors:** Adam Ciezarek, Reuben Gilbertson, Ellen Bell, David Murray, Eva Garnacho

## Abstract

Despite being two of the most commercially important flatfish (order Pleuronectiformes) in Europe, little is known of the population structure of common sole *Solea solea* and European plaice *Pleuronectes platessa*. To address this gap, we generated full-genome resequencing data for 244 sole and 189 plaice in the Celtic Sea and western English Channel region to analyse both neutral and adaptive loci and quantify population processes, such as reproductive isolation or adaptive differentiation in each species. For sole, there was no evidence of reproductive isolation or population structure at neutral loci. There was, however, adaptive differentiation as adaptive loci indicated two subpopulations, with separation in the western English Channel. This is consistent with previous studies using RAD-seq and gene-linked SNPs. For plaice, there was no evidence of population structure at either neutral or adaptive loci in the Celtic Seas and Western English Channel region. However, when considering a larger geographical area and utilising previously published genomic data, three distinct populations of plaice were identified (Iceland; North Sea, Kattegat and Western Baltic; Celtic Sea and western English Channel), with clear reproductive isolation indicated by neutral loci and adaptive differentiation indicated by adaptive loci. Moreover, three large chromosomal inversions were identified, which differed in their frequency between regions. These large structural variants represent putative key regions for adaptive differentiation. This study shows the benefit from quantifying neutral and adaptive loci to better understand population structure and genetic diversity of commercially important fish.

## Introduction

Common sole (hereafter sole), *Solea solea*, and European plaice (hereafter plaice) *Pleuronectes platessa*, are two of the most commercially important demersal flatfish in Europe (Jayasinghe et al., 2017). Sole is distributed from Scotland and southern Norway to Senegal, including the Mediterranean, North and western Baltic seas, whilst plaice is distributed from Portugal northwards to Iceland and the White Sea. Both species are caught in the same mixed demersal fishery in the Celtic Sea and western English channel (CWC) (Dunn & Pawson, 2002).

Despite the commercial importance of both species, relatively little is known of their population structure in the CWC. The CWC covers four ICES Divisions: 7.e (western English Channel), 7.f (Bristol Channel), 7.g (northern Celtic Sea) and 7.h (southern Celtic Sea; see Figure 1) Genomic techniques offer the potential to accurately quantify population differentiation and the contributions of genetic drift, selection and gene flow. Given the enormous population sizes of marine fish, and therefore likely reduced levels of genetic drift (Charlesworth, 2009), even a small amount of gene flow can result in very low levels of differentiation at neutral loci (loci showing no evidence of being under selection). As a result, adaptive loci (showing evidence of selection) have often been used to differentiate populations (Taylor et al., 2025). Full genomic sequencing is therefore important to identify population structure, as reduced representation approaches are likely to miss the small regions of the genome often involved with this adaptive differentiation (Andersson et al., 2024). In sole, two reduced-representation genomic approaches have been used to study population structure in the CWC. An earlier study based on 426 gene-linked SNPs (Single Nucleotide Polymorphisms) found significant differentiation at adaptive, but not neutral, loci between sole from the Celtic Sea/Bristol Channel and the western English Channel (Diopere et al., 2018). Furthermore, a RAD-seq analysis from 55,706 SNPs across the sole genome showed broad-scale population structure, with sole from the North Sea differentiated to those from the Bay of Biscay at neutral loci. At a finer scale, sole from south-western Ireland (ICES Division 7.j) were found to be distinct from sole across the Celtic Sea and western English Channel (Divisions 7.e-h) in neutral loci, and sole from the eastern part of 7.e were differentiated to those from the western part of 7.e and 7.f-h at adaptive loci (although not consistently between all statistical rectangles; Maes et al., 2025). No population genomic studies have been conducted to date across the region for plaice, with the whole genome sequencing and RAD-seq studies to date only carried out in Iceland, the North Sea, Kattegat and western Baltic (Le Moan et al., 2019, 2021; Weist et al., 2022). This work identified significant differentiation at neutral loci between plaice from Iceland to those from North Sea, Kattegat and western Baltic (hereafter NKB), with a pattern of isolation-by-distance identified across the NKB in adaptive loci.

**Figure 1.**
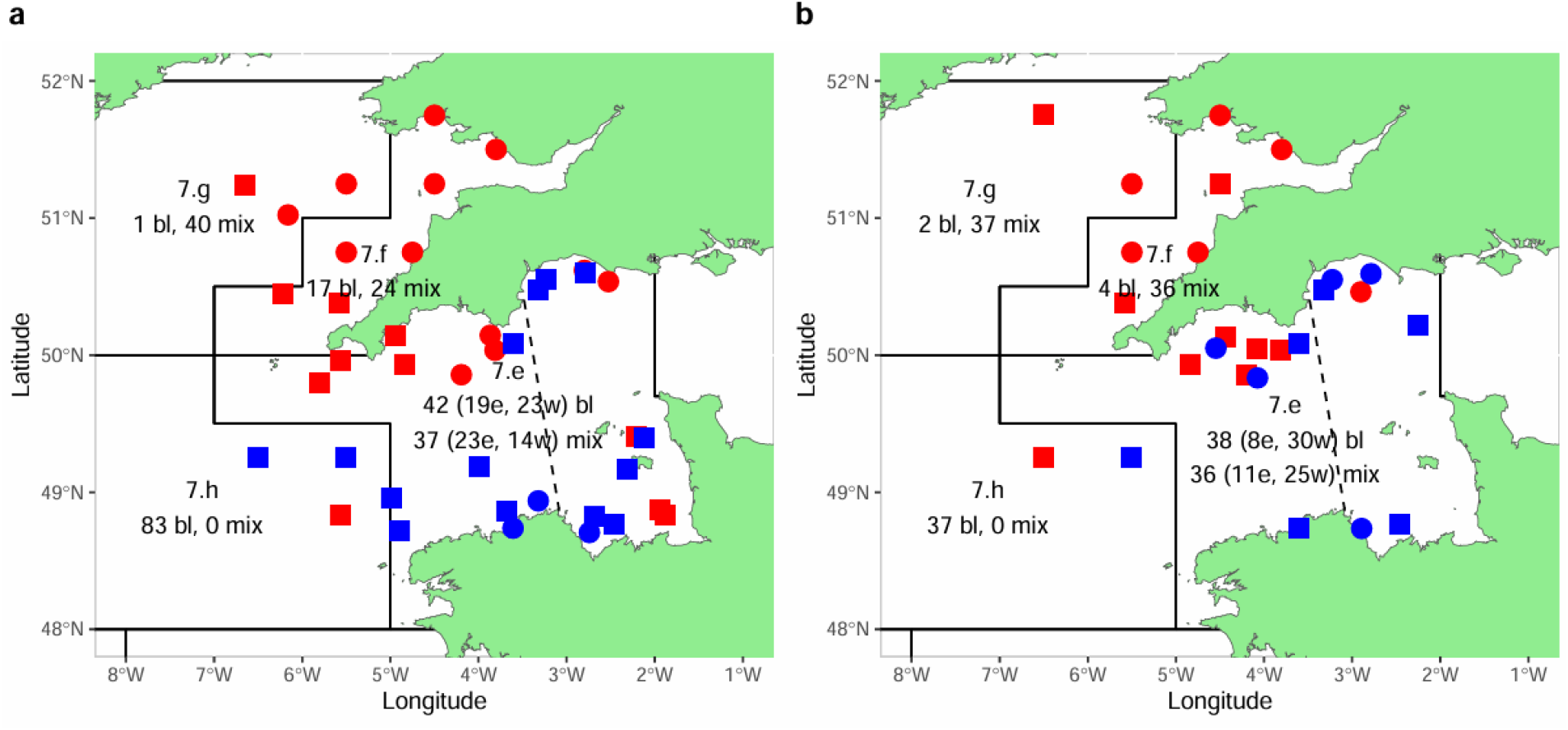
The number of baseline (bl) and mixed (mix) sole (a) and plaice (b) samples collected per ICES Division. The dashed line in 7.e represents the split between eastern and western 7.e used in analysis. The number of baselines and mixed samples in east (e) and west (w) are given. Squares represent sites where baseline samples were collected, circles where only mixed samples were collected. Points in red were sampled in 2023, with those in blue sampled in 2024.

Large structural variants, particularly chromosomal inversions, can act as drivers of divergence and local adaptation in the face of ongoing gene flow (Akopyan et al., 2025). Due to recombination suppression in inversion heterokaryotypes (Kirkpatrick, 2010), multiple adaptive loci may be inherited together, which may cause them to act as “supergenes” underpinning local adaptation. This has been documented in Atlantic cod, *Gadus morhua*, where such inversion supergenes are associated with different migratory behaviours (Kirubakaran et al., 2016; Matschiner et al., 2022). Conversely, inversions can accumulate recessive deleterious mutations at a greater rate than the collinear genome (Huang et al., 2022; Jay et al., 2021), thereby increasing the risk of inbreeding depression. Due to this importance for population divergence as well as adaptive and deleterious diversity within species, inversions must be accounted for to enable optimal management of genetic resources (Schneller et al., 2025). Chromosomal inversions have been identified in several marine fish, including plaice (Le Moan et al., 2019, 2021; Weist et al., 2022), but without any clear geographical structure in their presence or absence across the area studied (NKB and Iceland). To date, no work has identified chromosomal inversions in sole.

Here, we present whole-genome resequencing data for 244 sole and 190 plaice across the CWC (Divisions 7.e-h), supplemented with previously published data for 60 plaice individuals from the NKB and Iceland (ICES Division 5.a). Population structure and the within- and between-population differentiation at neutral and adaptive loci, including large chromosomal inversions, are investigated in both species separately.

## Materials and Methods

### Sampling and Sequencing

Sole and plaice samples from ICES Divisions 7.e-g were collected on [[removed for anonymity]] in January 2023 and January 2024, at the onset or just prior to spawning season for both species (Table S1). Samples from Division 7.h were collected by [[removed for anonymity]]. Muscle tissues were collected using the LVL rapid genetic sampling tool (LVL Technologies, Crailsheim, Germany) and stored in ethanol. Samples were classified as ‘baseline’ (meaning spawning individuals) or ‘mixed’ (unknown spawning status) based on sampling location, time and a sexual maturity score of at least 4 (ICES 5-stage maturity scale; ICES, 2023). Previously published short-read resequencing data for plaice in NKB was also downloaded from NCBI (Bioproject PRJEB36564; Weist et al., 2022).

DNA was extracted using the Maxwell® RSC Tissue DNA kit (Maxwell Technologies, San Diego, California). DNA quality control was carried out using gel electrophoresis and the Agilent 5400 fragment analyser system (Agilent Technologies, Santa Clara, California). Libraries were then prepared using the NovoGene NGS DNA library prep set and sequenced on the Illumina NovaSeq Xplus with 150bp paired-end reads (Novogene Corporation, Cambridge, England).

### Read mapping and SNP calling

Quality control of raw reads was carried out using fastp (v0.23.4; Chen et al., 2018), alongside trimming of adapter sequences and poly-G tails. Trimmed reads were mapped against their respective chromosome-level reference genomes (GCF_958295425.1 for sole, GCF_947347685.1 for plaice) using bwa mem (0.7.18; Li & Durbin, 2009), with resulting bam files sorted using samtools (v1.20; Li et al., 2009) and duplicates removed using the samremovedup.py script from GenErode (Kutschera et al., 2022). When sequence data from samples were split across multiple lanes, data from each lane were treated and mapped separately. Each sample was then genotyped individually, including both invariant and variant sites. Bcftools (v1.15.1) mpileup was used, filtering low quality alignments (Q < 30) and bases (Q < 30). SNPs were then called using the Bcftools consensus caller, filtering sites with high mapping depth (>2x genome-wide average), low mapping (depth < ⅓genome-wide average), low genotype quality (QUAL OR GQ < 30), or sites within 3 bases of an indel. A full variant and invariant dataset was then generated, merging data from all samples, retaining only sites with less than half missing data. A biallelic SNP set was also generated, retaining only biallelic SNPs with a minor-allele count of at least 3 and less than half missing data. Neighbour-joining trees were generated using VCF-kit (v0.2.6; Cook & Andersen, 2017) and inspected for any samples with long branch lengths, which were removed from subsequent analyses.

### Population genomic analyses

Differentiation across neutral and adaptive loci were used to classify population structure, using definitions adapted from Cadrin et al., (2014). Significant differentiation at neutral loci likely reflects reproductively distinct populations. A lack of differentiation at neutral loci with significant differentiation at adaptive loci likely reflects subpopulations, which have some degree of independence as well as connectivity to other subpopulations within a metapopulation.

Genome-wide neutral population genomic analyses were carried out separately for each species. Following filtering for linkage disequilibrium (R^2^ > 0.6 over 20kb windows), missing data (sites with > 20% removed) and minor allele-frequency (< 5% removed), up to ten rounds of PCA analysis were carried out using plink (v1.90; Purcell et al., 2007). With each round, outlier samples with a PC component at least six standard deviations from the mean were removed, until no outliers were left. Hierarchical clustering analyses were carried out on this filtered dataset with K = 1-5 carried out using ADMIXTURE (v 1.3.0; Alexander et al., 2009), with Cross-Validation (CV) error and the R package stockR (Foster, 2018), with AIC and BIC, to assess the optimal value of K. Global Hudson’s F_ST_, with standard error calculated using a block size of 1,000,000 bp, was also calculated between each pair of ICES Divisions using plink (v2.0.0; Chang et al., 2015). A 95% confidence interval was given by the mean F_ST_ ± 1.96 times the standard error. Any putative adaptive regions (F_ST_ Z ≥ 3 20kb windows, inversions or PBSN outlier windows, see below) were removed prior to F_ST_ computation. Significance was assessed by comparing against 2,000 random permutations (random reshuffling of ICES Division labels, counting matches where the lower bound of the 95% confidence interval exceeds the upper bound of the permutation 95% confidence interval), with Bonferroni corrected *p* < 0.01 used to assess significance. Given the adaptive differentiation previously identified between the eastern and western parts of 7.e (Maes et al., 2025), Division 7.e was analysed both as a whole and after dividing into eastern and western regions.

To estimate linkage disequilibrium (LD) and identify potential chromosomal inversions, pairwise R^2^ was calculated. Input SNPs were thinned to only include one SNP per kb, after removing sites with more than 10% missing data. Pairwise R^2^ for each SNP pair was calculated using vcftools (v0.1.16; Danecek et al., 2011), and emerald2windowldcounts.pl (https://github.com/owensgl/reformat, https://github.com/owensgl/haploblocks) was used to calculate the mean R^2^ between pairwise 100kb windows. Long blocks of elevated R^2^ were used to identify putative inversions. As the ends of chromosomes typically show elevated rates of recombination (Akopyan et al., 2022; Johnsson et al., 2021), small R^2^ peaks in these regions were not considered putative inversions. Furthermore, SuperInfer (https://github.com/PaulYannJay/SuperInfer), was used to carry out local PCA, identifying regions with high clustering scores at K=2 or K=3 as well as with high distance between centroids of the most distant clusters at K=3, typical of inversions. Input vcfs were phased using shapeit (v2.r904), after removing sites or samples with ≥ 10% missing data. SuperInfer was then run on 50kb windows, with 10kb overlap and clustering on five principal components. Inversions were identified when there were clear regions with elevated R^2^, high clustering scores at K=2 or K=3 and a high cluster distance. Inversions were further validated if individuals with the heterozygotic AB genotype at the inversion (identified using PCA), had significantly elevated heterozygosity compared to AA or BB individuals.

### Identification of adaptive loci

In order to identify whether any Division had a significant excess of high F_ST_ SNPs (Hudson’s F_ST_ ≥ 0.3) compared to other Divisions, Hudson’s F_ST_ for each SNP between each pair of ICES Divisions were calculated using piawka (v0.8.8; https://github.com/novikovalab/piawka). The count of high F_ST_ SNPs was compared against 100 random permutations, whereby ICES Division labels were randomly shuffled. A complete lack of overlap (*p* = 0) between the observed and permutation high F_ST_ SNP counts was used to identify putative subpopulations with potential adaptive marker differentiation between them. These were further validated by tests for SNPs under selection.

In cases with exactly two putative subpopulations, or where neutral loci indicated genetic differentiation between exactly two groups (the *npop* = 2 case), two further tests were carried out to identify if there regions or SNPs showing evidence of positive selection (adaptive loci): i) Individual SNPs with evidence of selection were identified using Bayescan (v2.1; q ≤ 0.05, pr_odds 10000), and ii) 20kb windows with evidence of selective sweeps were identified. These were defined as windows with elevated Hudson’s F_ST_ (Z ≥ 3) and either the lowest 5% of log2-transformed π1/π2 or π2/π1 ratios, whereby π1 and π2 represent π in either tested ICES Division, or the lowest 5% of Tajima’s D values for either Division. Piawka was used to calculated Hudson’s F_ST_ per 20kb window, with pixy (v1.2.7) (Korunes & Samuk, 2021) used to calculate π and D_XY_ and VCF-kit used to calculate Tajima’s D. Transformations (Z and log2) were carried out using custom python scripts (available on github upon publication), requiring at least 200 SNPs per window for F_ST_ or Tajima’s D or 5,000 sites per window for π or D_XY_.

In cases with at least three putative subpopulations, or where neutral loci indicated genetic differentiation between at least three groups (the *npop* ≥ 3 case), the normalised Population Branch Statistic (PBSN) was used to identify putatively adaptive regions (Malaspinas et al., 2016; Shpak et al., 2025). This converts pairwise F_ST_ estimates between three groups of individuals to branch lengths for each group, normalised by outgroup branch lengths at the same genomic region. Piawka was run calculating Hudson’s F_ST_ per 20kb window. PBSN was then calculated on all windows with at least 200 SNPs using custom python scripts (available on github upon publication). F_ST_ values were converted to a branch length (T) using the equation T = -log(1-F_ST_), with PBSN calculated for each ICES Division

(a, b and c in turn) using the equation 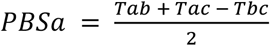 and then 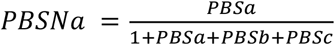. PBSN values were plotted across the genome, with outliers (Z ≥ 3) highlighted. These PBSN outliers reflect regions where one ICES Division is significantly differentiated from two other tested Divisions, normalised by the total differentiation existing across them.

As increased F_ST_ can be driven by either increased D_XY_ or decreased π 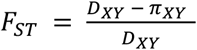, where 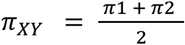, values of D_XY_ and π_XY_ in outlier windows were compared against values outside of putative selective sweeps. High F_ST_ in combination with increased D_XY_ reflects accumulated genetic variation built up by mutations since two populations split, whereas high F_ST_ in combination with decreased π reflects loss of diversity in small populations or selective sweeps (de Jong et al., 2025). High F_ST_ in combination with both low D_XY_ and low π would likely reflect reduced ancestral diversity, possibly as a result of recurrent selective sweeps (in either population as well as their common ancestor), linked selection or demographic events (Chase et al., 2021). In the *npop* = 2 case outlined above, π_MAX_ (the π of the ICES Division with the highest π value in a window) was used as the measure of π, as ratios of π were used to identify selective sweeps (which would therefore result in reduced π_XY_ not independent of the test). In the *npop* ≥ 3 case above, π_XY_ was used (adapted as e.g. 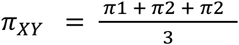for three populations). To assess significance, 10,000 sets of *n* 20kb windows (where *n* is the number of selective sweep 20kb windows found) were randomly selected and D_XY_ and π calculated for them, with *p* < 0.01 used to assess significantly increased or decreased D_XY_, π_XY_ or *π*_MAX_.

### Assignment models

If population structure was identified, mixed individuals were tested to see if they could be assigned to any of the reference populations. Three different subsets of SNPs were tested, each of which were filtered for linkage disequilibrium: i) SNPs under selection (Bayescan *q* < 0.01 and in outlier window); ii) SNPs with F_ST_ > 0.3 and iii) Ancestry Informative SNPs with allele-frequency differences (AF) of at least 0.3 between all pairs of ICES Division in split (e.g. if p1 is differentiated from p2 and p3, but p2 and p3 are not differentiated, SNPs with AF > 0.3 between p1 and p2, but not between p2 and p3). These were tested for clear separation of baseline samples in PCA analysis. Assignments of mixed samples were made using the R Package assignPOP, utilising the Naive Bayes, lda and SVM radial and linear models (with gamma and SVM cost optimisation utilising the e1071 package). Input vcf files were converted to baseline and mixed genepop files using custom python scripts (available on github upon publication). Model fits were tested using Monte-Carlo and k-fold Cross-Validation to assess how well baselines were self-assigned. Assignments were also made using stockR, using mixture models followed by bootstrap resampling. For the stockR analysis, baseline samples of the larger population were randomly down sampled so that sample size was even between the tested populations.

## Results

### Sampling, read mapping and SNP calling

A total of 244 sole and 190 plaice were sampled and sequenced (Figure 1). Mapping depths ranged from 7-65x for sole and 4-44x for plaice. A total of 29.2M and 22.3M biallelic SNPs were called for the sole and plaice respectively, with 568Mb and 539Mb sites genotyped (including invariant sites). Following inspection of neighbour-joining trees and low mapping depths, a further four baseline sole samples were removed from the dataset (one from eastern 7.e and three from 7.h) alongside one baseline plaice from western 7.e and two mixed plaice (both from 7.g). The number of baseline sole and plaice from each unique sampling station are given in Table S2. After adding the 63 plaice from PRJEB3656, a total of 23.9 SNPs were called from 250 plaice individuals.

### Sole population structure

Genome-wide PCAs for sole showed no signal of population structure, with ADMIXTURE and stockR identifying K=1 as the optimal number of populations (Figure 2a). Following the removal of high-F_ST_ regions (see below), neutral F_ST_ values between sole from different ICES Divisions were very small with no evidence of differentiation (F_ST_ < 0.0006; adjusted *p* > 0.1; Table 1). ICES Division 7.g was excluded from comparisons due to the lack of baseline samples (n=1). There were no candidate inversions identified by R^2^ hotspots (all pdfs available on dryad upon publication). The only regions with elevated R^2^ were very small as well as being either in regions with missing SNPs or near the start or end of chromosomes.

**Figure 2.**
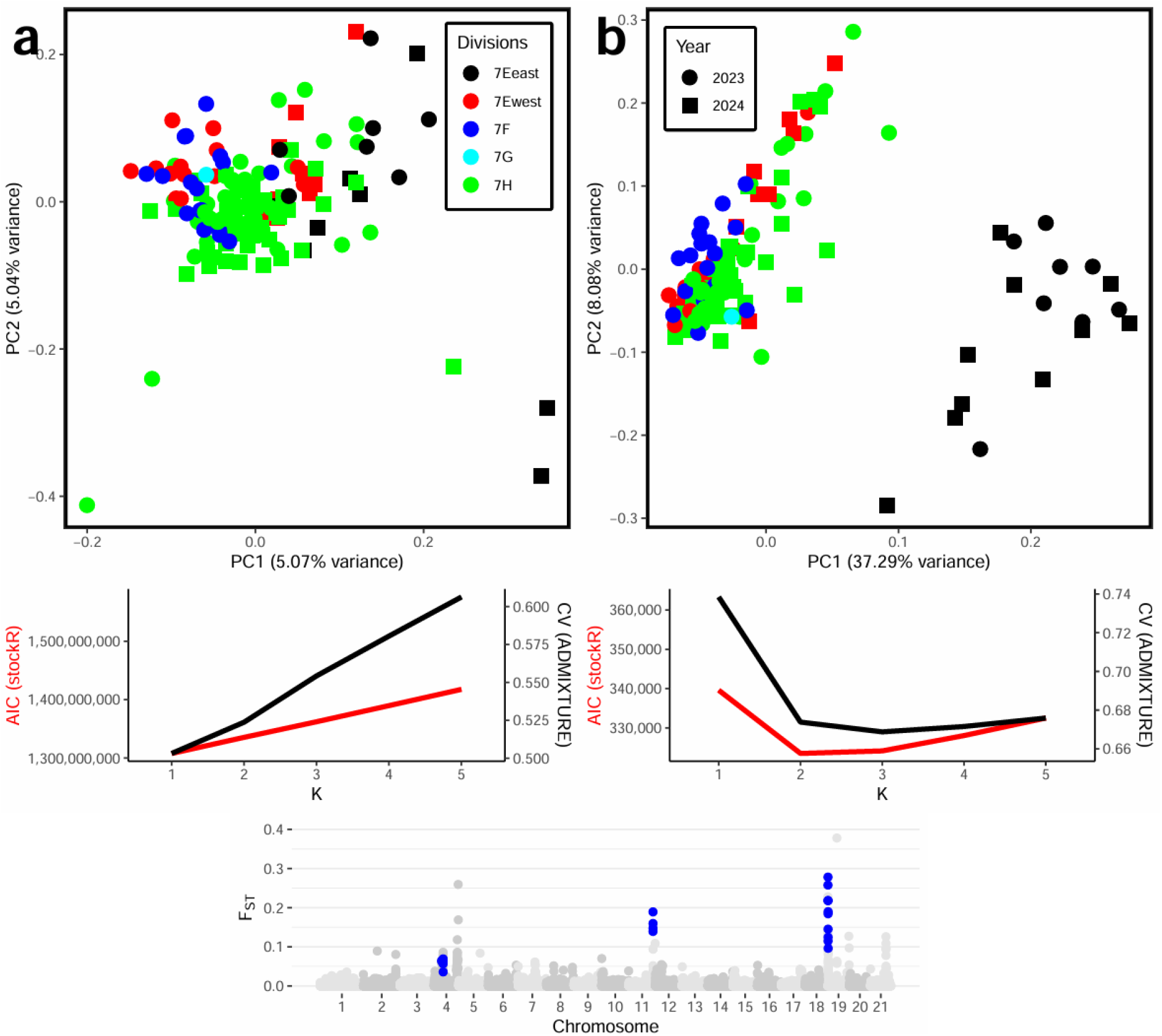
PCA plots for sole, with Admixture Cross-Validation and stockR AIC at K=1-5 populations for a) 7.2M SNPs across the genome; and b) PCA of 1,457 Ancestry Informative SNPs between eastern 7.e and western 7.e, 7.f-h sole. The legends in panels A and B apply to both plots. c) F_ST_ between eastern 7.e and western 7.e, 7.f-h sole, plotted in 20kb windows. Windows containing the 483 adaptive loci are highlighted in blue.

**Table 1.** Mean neutral F_ST_ between baseline sole from each ICES Division. Bonferroni corrected *p* values for each comparison against 2,000 random permutations.

|  | 7.e (west) | 7.f | 7.h |
| --- | --- | --- | --- |
| 7.e (east) | 0.0006 ($p = 0.3$) | 0.0006 ($p = 0.2$) | 0.0005 ($p = 0.1$) |
| 7.e (west) | | 0.0004 ($p = 1$) | 0.0003 ($p = 0.5$) |
| 7.f | | | 0.0003 ($p = 0.4$) |
| 7.e (whole) | | 0.0003 ($p = 0.2$) | 0.0002 ($p = 0.2$) |

A significant excess of high F_ST_ SNPs was consistently found between sole from eastern 7.e and all other ICES Divisions, including western 7.e (Table S3; *p* = 0). No significant excess was found between any other ICES Divisions (7.e west, 7,e as a whole, 7.f or 7.h). Bayescan identified 223 SNPs to be under selection between the two groups (eastern 7.e and western 7.e,f-h) and 154 20kb windows were found with elevated F_ST_ and either reduced Tajima’s D or decreased π in either the western 7.e + 7.f-h group (99) or the eastern 7.e group (63). Both D_XY_ (0.004 within compared to 0.006 outside; *p* = 0) and π_MAX_ (0.005 within and 0.006 outside; *p*=0) were reduced inside the 154 20kb windows compared to the rest of the genome. 210 SNPs of the Bayescan q ≤ 0.05 SNPs were found in 20 of the 154kb outlier windows (13 SNPs within one 1.6Mb region of chromosome 4, 61 within one 8-kb region of chromosome 11, and 136 one 220kb region of chromosome 19) (Figure 2c). PCA analysis on 45 of these SNPs (following filtering for linkage disequilibrium) did not reveal clear population structure, likely due to the low number of SNPs (Figure S1).

### Sole assignment model

Assignment models were tested on three datasets, each using all the baseline eastern 7.e individuals as reference for one population and all other baselines as reference for the other: i) 45 SNPs under selection (with a lighter filter of R^2^ >= 0.8 instead of 0.6 due to the low number of SNPs; ii) 127 high F_ST_ SNPs; iii) 1,457 Ancestry Informative SNPs. Datasets i) and ii) did not show complete segregation of the populations in the PCA (Figure S1) and could not self-assign baseline eastern 7.e samples in PopAssign kfold analysis. Dataset iii) showed distinct separation of baseline samples in the PCA (Figure 2b), and both k-fold and MC showed each tested PopAssign model could consistently (at least 17/18 baselines correctly assigned) self-assign baselines. However, results between PopAssign models were highly inconsistent (Table S4). Results were also inconsistent with stockR assignments, which showed a lack of power in discriminating the eastern 7.e population, with 32 individuals (including the 18 baselines) identified as most likely to belong to the eastern 7.e subpopulation, all except one of which were found in either eastern or southern 7.e (Table S4). However, bootstrapping analysis revealed only 20/34 had at least an assignment probability > 0.95 of belonging to this population, with the remaining 14 unassigned (assignment probability < 0.95), including all mixed samples. 79 of the 88 samples assigned by stockR as belonging to the western 7.e + 7.f-h subpopulation had assignment probability

> 0.95, Results of PopAssign and stockR models were consistent for all mixed individuals outside of 7.e, with all consistently identified as belonging to the western 7.e + 7.f-h subpopulation.

### Plaice population structure

PCA on the full genome-wide data showed a pattern broadly consistent with inversions, with three groups along PC1 (Figure 3a). When inversions were removed, no population structure was apparent, with stockR and ADMIXTURE both having best support for K=1 populations (Figure S2). F_ST_ values in these regions between plaice from different ICES Divisions were very small, with no evidence of significant differentiation (F_ST_ < 0.001, all *p* > 0.4, Table 2). Due to the low number of baseline samples, F_ST_ was not calculated in ICES Divisions 7.f (n = 4) or 7.g (n = 2). There was also no excess of high F_ST_ single SNPs between any ICES Division (All Bonferroni adjusted *p* > 0.07; Table S5).

**Figure 3.**
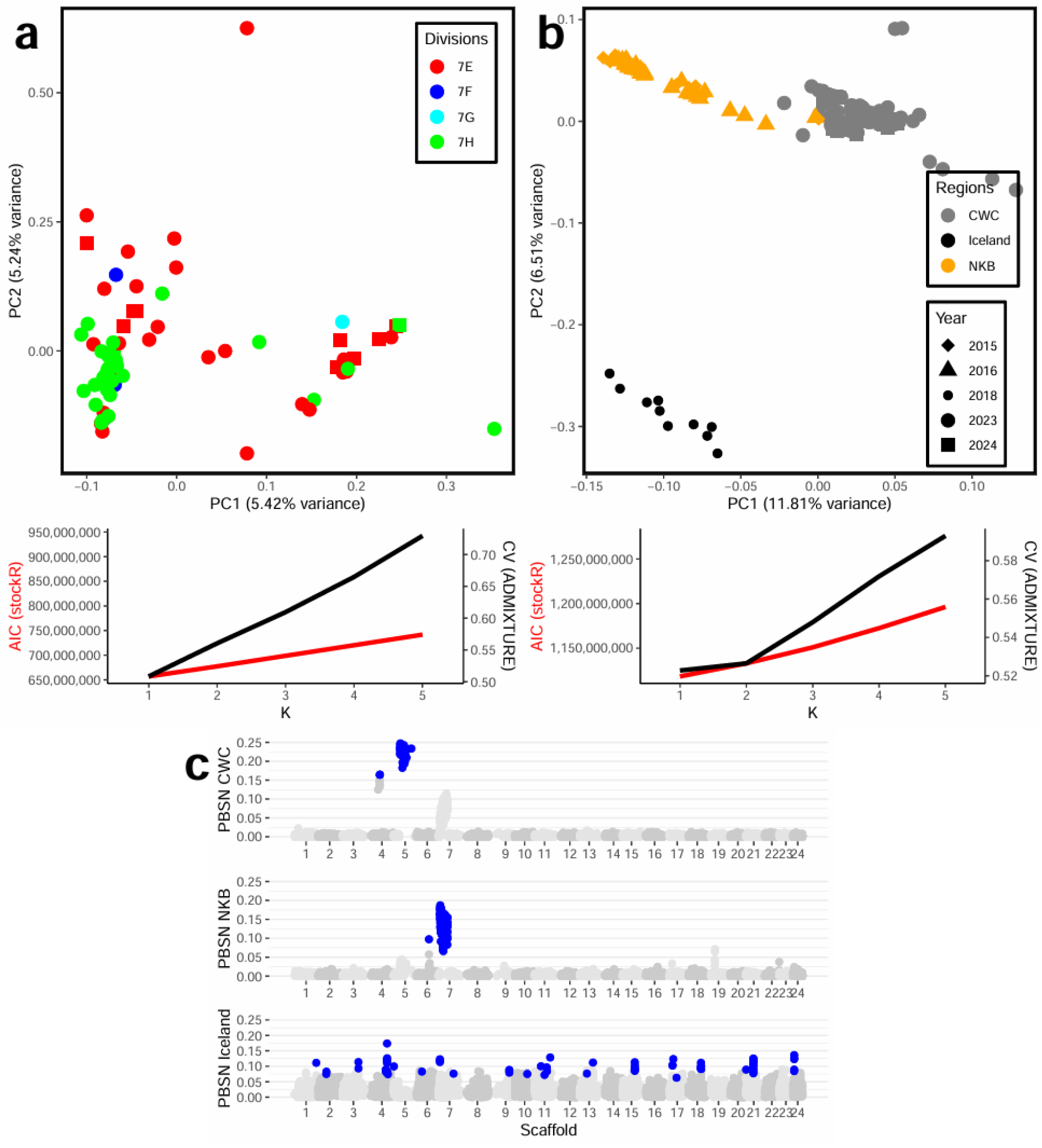
PCA plots for plaice, with Admixture Cross-Validation and stockR AIC at K=1-5 populations for a) 6M SNPs across the genome for the Celtic Seas and western English Channel (CWC) samples collected in this study; b) 6M SNPs for the same samples in addition to those from North Sea, Kattegat and western Baltic (NKB) and Iceland (the year legend in panel b applies to both plots); and c) PBSN statistics calculated for the CWC, NKB and Iceland populations, with outlier (Z ≥ 3) values highlighted in blue.

**Table 2.** Mean neutral F_ST_ (excluding all inversions or PBSN outliers) for plaice, between baseline samples from each ICES Division, as well as at the larger regional scale between CWC (Celtic Seas and western English Channel). NKB (North Sea, Kattegat and western Baltic) and Iceland. Bonferroni corrected *p* values for each comparison against 2,000 random permutations.

|  | 7.e (west) | 7.h | NKB | Iceland |
| --- | --- | --- | --- | --- |
| 7.e (east) | 0.0002 ($p = 1$) | 0.001 ($p = 1$) | | |
| 7.e (west) | | 0.0002 ($p = 1$) | | |
| 7.e (whole) | | 0.0003 ($p = 0.4$) | | |
| CWC | | | 0.005 ($p = 0$) | 0.03 ($p = 0$) |
| NKB | | | | 0.03 ($p = 0$) |

When data from Iceland and NKB were included (alongside the mixed CWC samples, due to the lack of population structure within CWC), three populations were apparent in the PCA, although stockR and ADMIXTURE both still had best fits at K=1 (Figure 3b). Neutral F_ST_ was significantly elevated in all comparisons between Iceland, NKB and the CWC (Table 2, all *p* = 0). Patterns of R^2^, alongside a high cluster distance and PCA peaks for K=2 or K=3 (Figure S3, Figure S4) indicated inversions in chromosomes 4, 5 and 7. These were supported by elevated heterozygosity in the AB karyotype. Individuals in the AA karyotype consistently had elevated heterozygosity compared to the BB karyotype (*p=*0). This was likely due to over-representation of the Iceland and NKB samples in the AA karyotype, which had reduced heterozygosity (average=0.01) compared to the Celtic Seas samples (average=0.02, t-test *p*=0; Figure S5). Within the CWC, no population structure or geographical pattern was apparent either within PCAs, or when individuals with each karyotype were mapped (Figure 4). However, there was differentiation between the NKB and Iceland and the CWC. AA karyotypes for each inversion were found predominantly in Iceland and NKB, with BB dominating in CWC. The chromosomes 4 and 5 AB karyotypes were found predominantly in CWC, whereas the chromosome 7 AB karyotype was predominantly in Iceland and NKB. PBSN outliers for CWC were exclusively found in these three inversions, with NKB outliers only in the chromosome 7 inversion. By contrast, 274 PBSN outliers for Iceland were distributed across the genome (Figure 3c). These PBSN outliers for Iceland had significantly reduced D_XY_ (D_XY_ 0.006 against NKB inside outlier windows, 0.008 outside; *p*=0, D_XY_ 0.007 against CWC inside outlier windows, 0.008 outside; *p*=0) as well as π_XY_ (0.005 inside outlier windows, 0.007 outside; *p*=0).

**Figure 4.**
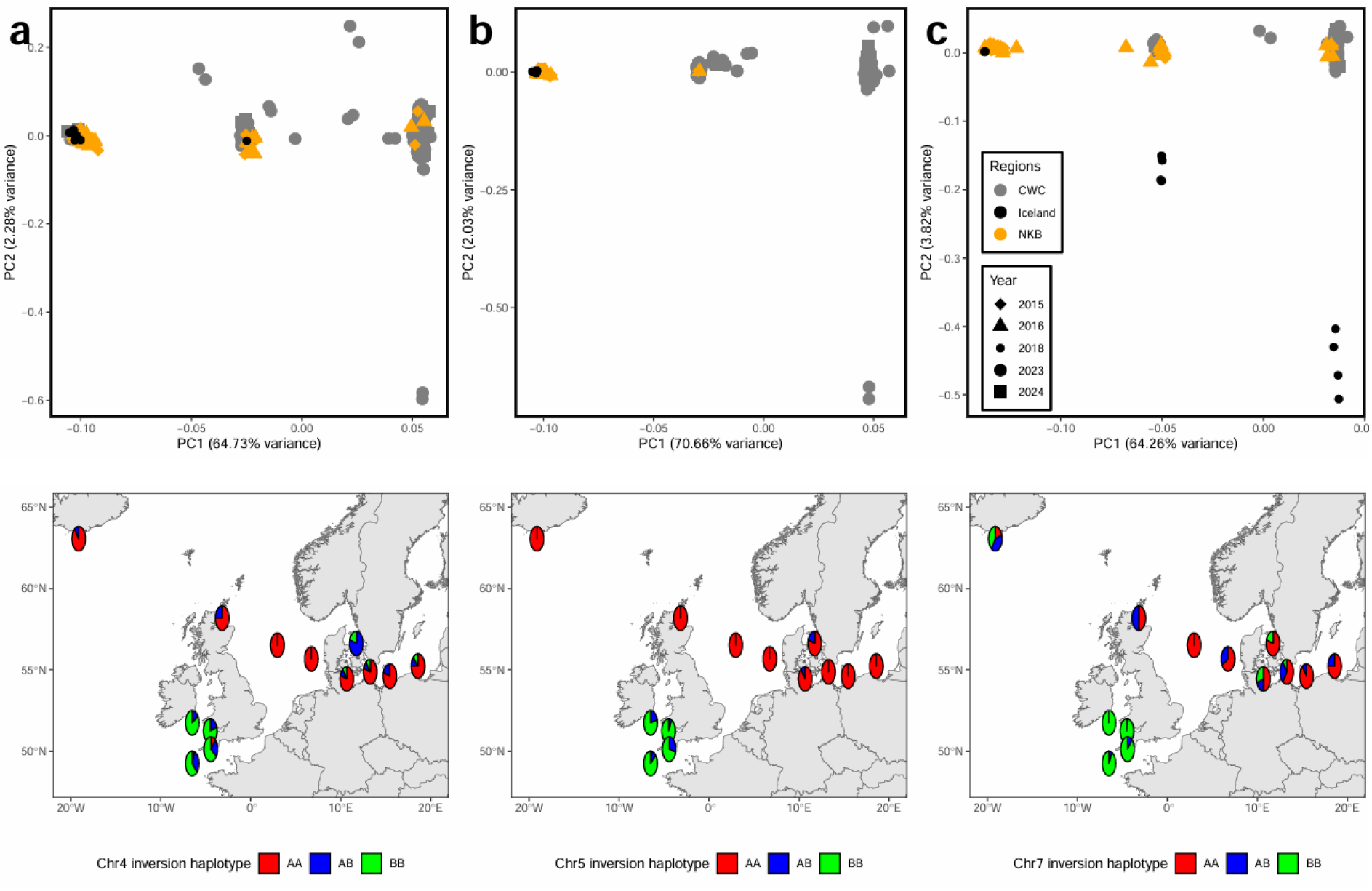
Inversion haplotypes of plaice samples for the inversion at chromosome 4 (a), chromosome 5 (b) and chromosome 7 (c). The top row shows PCA for all SNPs in each inversion region, each showing three clusters (AA on left, AB in middle and BB on right). The bottom row shows the frequency of each haplotype per sampling location. For the CWC samples, samples were aggregated per ICES Division for clarity.

## Discussion

Our results show no reproductive isolation for either the plaice or sole across the CWC as indicated by neutral loci. We find evidence of adaptive differentiation for sole in putative adaptive loci, with sole from the east of ICES Division 7.e to be distinct to those from western 7.e as well as 7.f-h, indicating two subpopulations within a metapopulation. In contrast, there was no evidence of adaptive differentiation for plaice as indicated by adaptive loci in the CWC. There was, however, evidence for different populations when including NKB and Iceland, as indicated by neutral loci differentiation and adaptive loci differentiation at large chromosomal inversions.

Our findings build on previous studies in demonstrating adaptive differentiation between sole from eastern and western 7.e. Although Diopere et al., (2018) found adaptive differentiation in their gene-linked SNPs between 7.e and 7.f, their sampling only included the eastern, but not western section of 7.e. Our study showed that the differentiation they found was between 7.f and eastern 7.e specifically, rather than 7.e as a whole. A recent RAD-seq study also found significant differentiation at adaptive loci between all except one statistical rectangle in western 7.e or 7.f-h and the eastern section of 7.e (Maes et al., 2025). The adaptive loci we identified largely overlapped with those identified by RAD-seq, with clusters in the same regions of chromosomes 4,11 and 19 (Sarah Maes personal communication). This consistency between results across three studies with differing sampling schemes, genomic data types and methods of identifying adaptive loci provides compelling evidence for adaptive differentiation in sole between eastern and western 7.e, with western 7.e undifferentiated from 7.f-h. This would be consistent with the two groups representing distinct but connected subpopulations, within a larger metapopulation (Atmore et al., 2024; Cadrin et al., 2014). A population split either side of the Hurd Deep, located north of the Channel Islands, has also been proposed based on fisheries and tagging data showing movement of sole from 7d into eastern 7.e (Burt & Millner, 2008). Further baseline sampling will be necessary to identify the exact geographic barrier between eastern and western 7.e, as well as the full geographical extent of each subpopulation.

The adaptive regions of the genome underpinning the definition of this subpopulation also had significantly reduced π and D_XY_ compared to the genome-wide averages. This suggests low levels of ancestral variation, possibly as a result of ongoing and ancient selection recurrently acting on the same genomic regions (Chase et al., 2021). These low levels of differentiation may explain the lack of power when these loci are used to assign mixed individuals to either subpopulation, evidenced by the inconsistent results from assignment models. It is possible that loci will have more power in cases with elevated D_XY_ alongside F_ST_, indicating an accumulation of mutations accumulating following a population split, rather than selection acting on low levels of standing variation (de Jong et al., 2025) as we observe here. This may also have been influenced by the skewed baseline population sizes (Birchard et al., 2025), with only 18 baselines from the eastern 7.e subpopulation compared to 118 from the western. Further mixed and baseline sampling may be necessary to produce mixing maps within the region and refine the optimal SNP set for assignment models.

Our study presents the first population genomic data for plaice in the Celtic Sea and western English Channel (CWC) region, with no evidence of any population structure in either neutral or adaptive loci. This is consistent with tagging data for plaice, which has shown that plaice from ICES Divisions 7.a and 7.e-f mix within Division 7.g (Dunn & Pawson, 2002). By incorporating previously published genomic data from Iceland and NKB, (Le Moan et al., 2019, 2021; Weist et al., 2022), we find that plaice in the CWC are genetically distinct to both the NKB group and the Icelandic group in both neutral and adaptive loci, likely reflecting three significantly reproductively isolated populations. However, further baseline sampling is necessary to identify where the boundaries between these populations lie. The split between Icelandic and NKB plaice is long established and was found by the original authors who published the genomic data (Weist et al., 2022) as well as previous studies using microsatellites and mitochondrial DNA (Hoarau et al., 2004; Was et al., 2010). We cannot rule out a possible batch effect as the datasets were collected and sequenced at different times and using different technologies. We did observe higher heterozygosity in the Celtic Sea plaice, likely as a result of higher sequencing depth, which would support this.

However, we would not expect a batch effect to result in the strong signals associated with the specific inversions we observed, or for the CWC plaice to be more similar to the NKB than Iceland if the differentiation was driven by batch-effect alone. We note that it may slightly elevate patterns of neutral divergence across the genome. Plaice represents an interesting species to further test the complex relationships between local adaptation and the adaptive and deleterious variation found on inversions (Schneller et al., 2025) across their wide geographical distribution and across time (e.g. in relation to environmental change).

## Conclusions

Expanding baseline sampling of plaice and sole will be important to investigate the extent of each identified subpopulation. In particular, sampling in the southern North Sea (ICES Division 4.c) and Eastern English Channel (7.d) will be important to quantify mixing between the North Sea and the CWC. Given the increasing trends and projected regional changes in sea temperature around the UK (Cornes et al., 2025), the possibility of subpopulations and metapopulation changes in fish populations (van Nouhuys, 2009), and other forecasts on habitats and fish distribution changes (Townhill et al., 2023), it is important to further investigate the genetic processes of sole and plaice using a temporal baseline. Furthermore, the Celtic Seas, English Channel and North Sea are important for a wide range of commercial fish species, and are home to a wide range of biogeographical features which may act as barriers to gene flow. Wide-ranging genomic studies of a diverse range of species across the region will support management of these valuable fishery resources.

Significant adaptive differentiation with very low or no neutral differentiation has been documented in a wide range of marine fish, at varying spatial scales and using different techniques for identifying adaptive or neutral loci (Fuentes-Pardo et al., 2023; Martinez Barrio et al., 2016; McKeown et al., 2020). This may reflect local adaptation even in the face of ongoing gene flow, which is likely due to high dispersal capacity and a lack of physical boundaries in the ocean. This would be consistent with complex metapopulation structures (Cecino & Treml, 2021). However, metapopulation dynamics are just one of many factors which may reduce neutral diversity in marine fish. Selective processes may reduce neutral diversity, as the enormous census population size of marine fish could mean highly efficient removal of even slightly deleterious (near-neutral) mutations, and therefore widespread selective sweeps removing linked sites (Charlesworth & Jensen, 2022). Further demographic processes such as population size changes and sweepstakes reproduction, which may be accompanied by rapid selective sweeps further reducing diversity (Eldon & Stephan, 2024) may also act to reduce diversity as could neutral processes such as mutational bias, biased gene conversion or a negative relationship between population size and mutation rate (Charlesworth & Jensen, 2022).

Given the importance of understanding the drivers of neutral and adaptive diversity in marine fish for effective management. This will require population genomics to be combined with fisheries and larval survey work to identify spawning grounds and collect accurate baseline samples for population designation, as well as tagging and otolith studies to quantify the movements of fish over their lifespan. In addition to spatial data, it is also critical to collect ecological data, as genetically distinct ecotypes may exist within the same regions (del Rio et al., 2025). Emerging genomic techniques such as Close-Kin Mark Recapture (CKMR; Bravington et al., 2016) and pangenomics (Secomandi et al., 2023) will also help to further understand the dynamics driving marine fish diversity. These will enable accurate quantification of census population sizes to compare genetic diversity against (CKMR), and accurate analysis of large structural variants, such as inversions as well as thorough quantification of adaptive and neutral diversity (pangenomics). We show here that population structure of commercially important marine fish may be driven by small fractions of the genome. This highlights the need to integrate these methods with emerging genomic technologies as well as different data sources to accurately identify population structure, thereby supporting effective fisheries management.

## Data availability

Raw sequencing reads will be made available on the European Nucleotide Archive upon publication. All custom python and R code as well as PCA output files will be available on Zenodo and Dryad respectively upon publication.

## Acknowledgements

We wish to thank the Fishing Industry partners at the South West Fisheries Alliance. We also thank Ian Holmes for the invaluable contribution from Cefas Fisheries data collection framework and colleagues taking samples for genetics at RV Cefas Endeavour, Matt Eade, Joseph Ribeiro, Kathrin Vossen, Chyanna Allison, Victoria Campon-Linares, Joseph Watson, Aaron Brazier, Susan Kenyon, Fiona Gilmour, and Scientists in Charge (SICs). We wish to thank Jim Ellis for his helpful review and comments on the manuscript. AC, RG, DM and EG were supported by Defra Project FRD050.

