## Supplementary Materials for "Genomics reveals population structure despite high connectivity of common sole, *Solea solea*, and European plaice, *Pleuronectes platessa*, in the Celtic Sea and western English Channel"

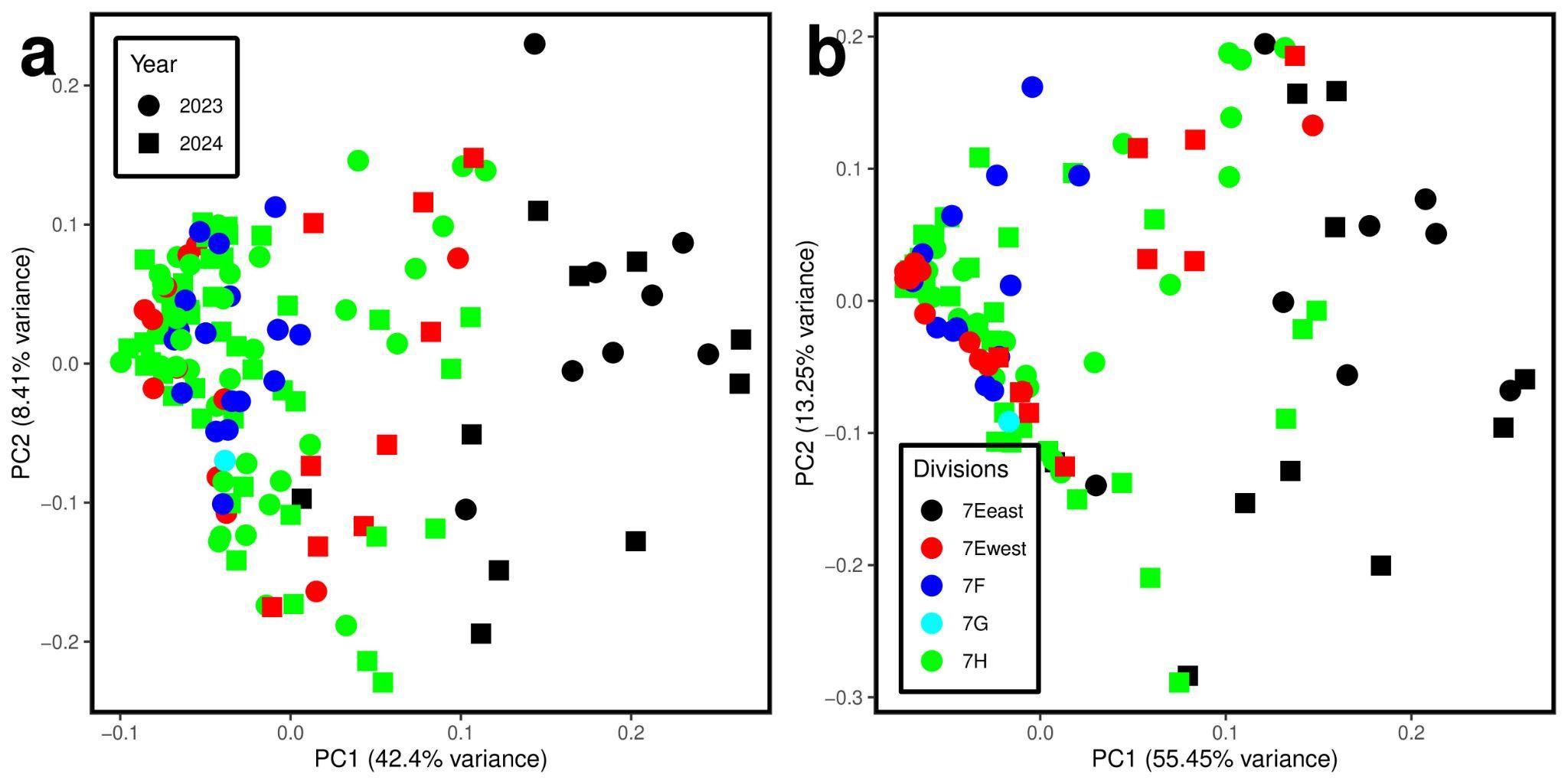

Figure S1. PCA analysis on baseline sole for a) 127 Fst ≥ 0.3 SNPs; and b) 45 SNPs under selection (Bayescan *q* < 0.05 and in an outlier window).

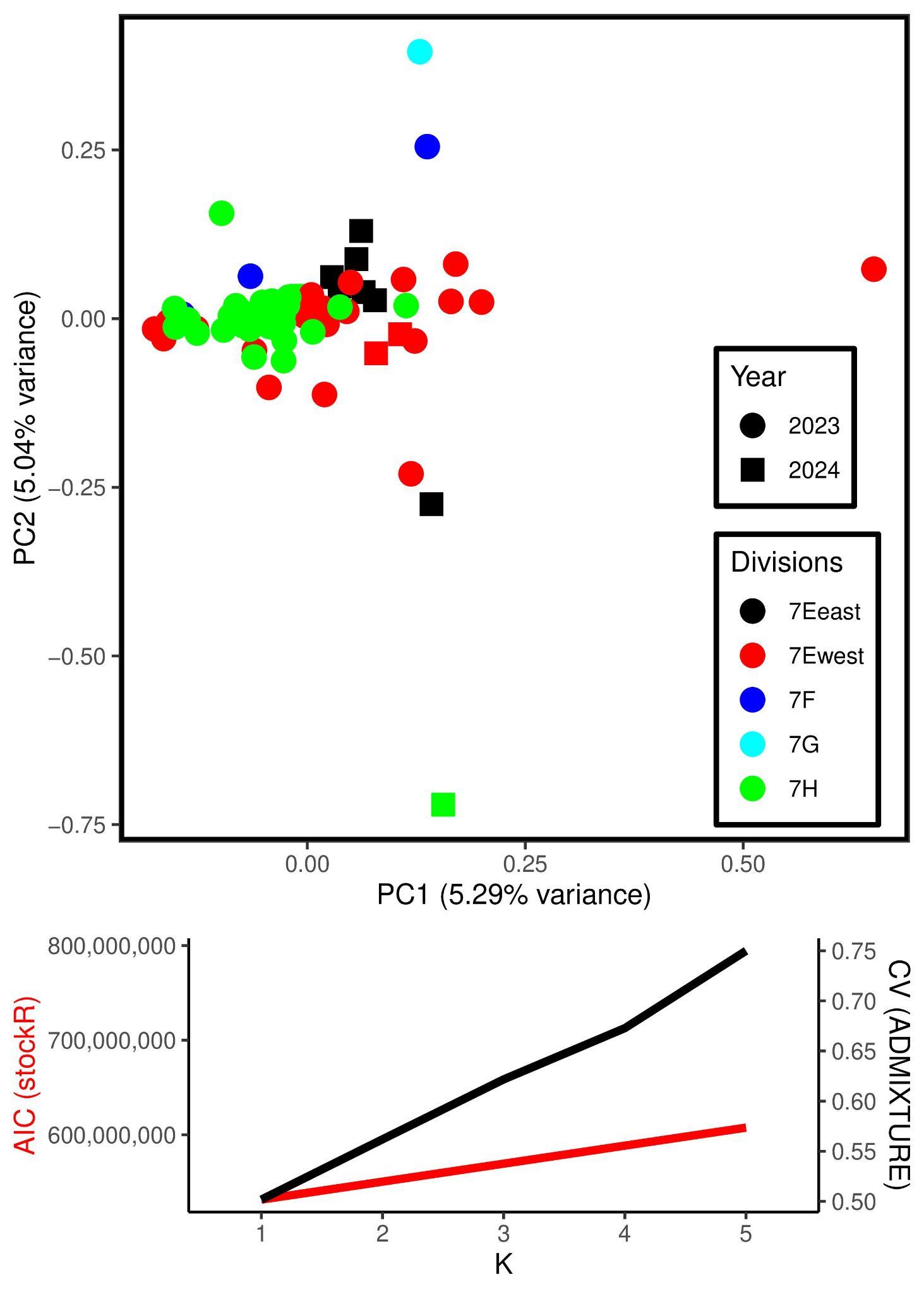

Figure S2. Plaice PCA plot, with Admixture Cross-Validation and stockR AIC at K=1-5 populations for a) 5.8M SNPs across the genome for the Celtic Seas and Western Channel (CWC) samples collected in this study, excluding inversions on chromosomes 4, 5 and 7

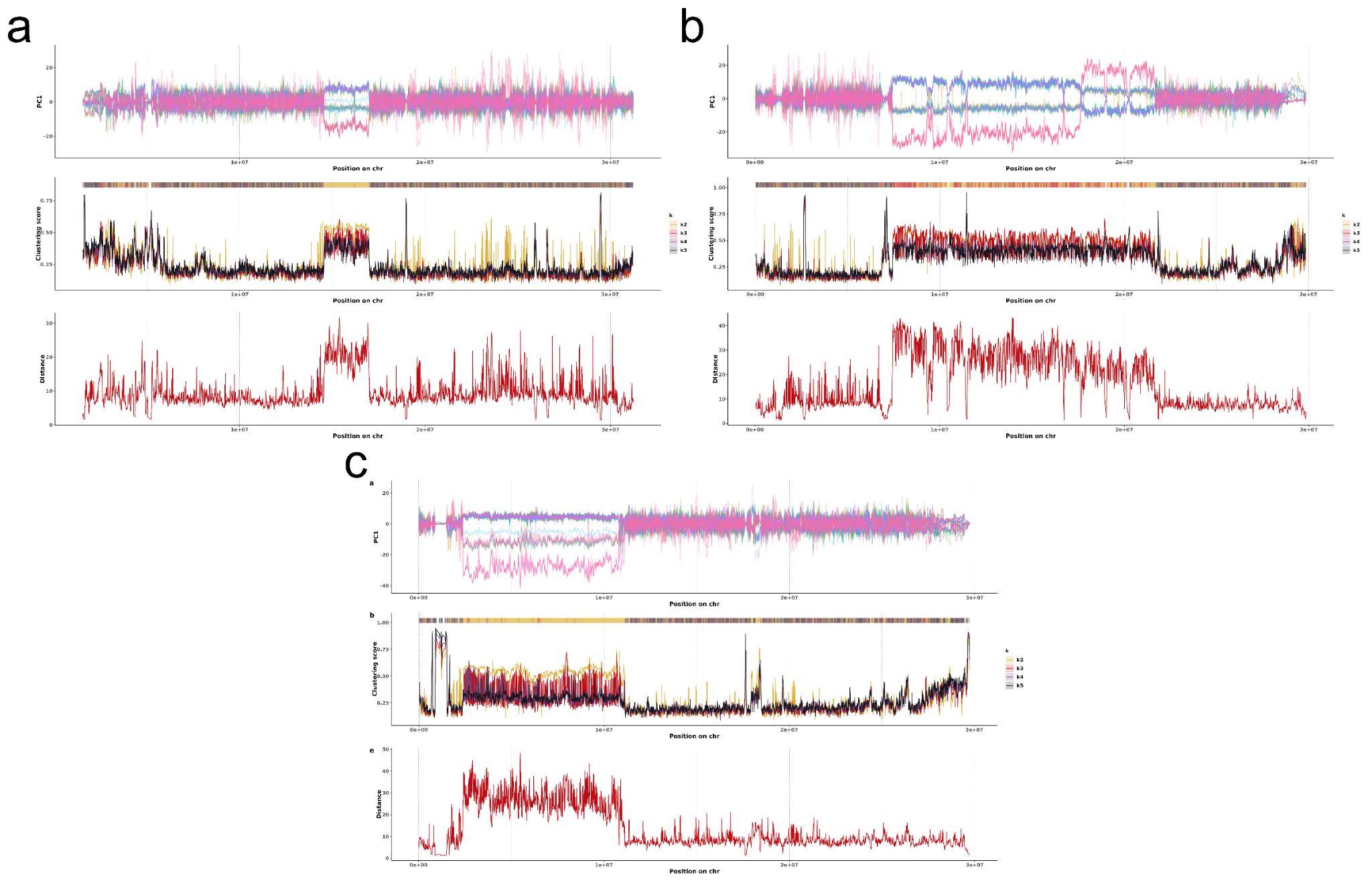

Figure S3. SuperInfer output for plaice chromosomes 4 (a), 5 (b) and 7 (c). Top row shows PC1 score, second row shows clustering score at K=2-5 and bottom row shows distance between centroids of the most distant clusters at K=3.

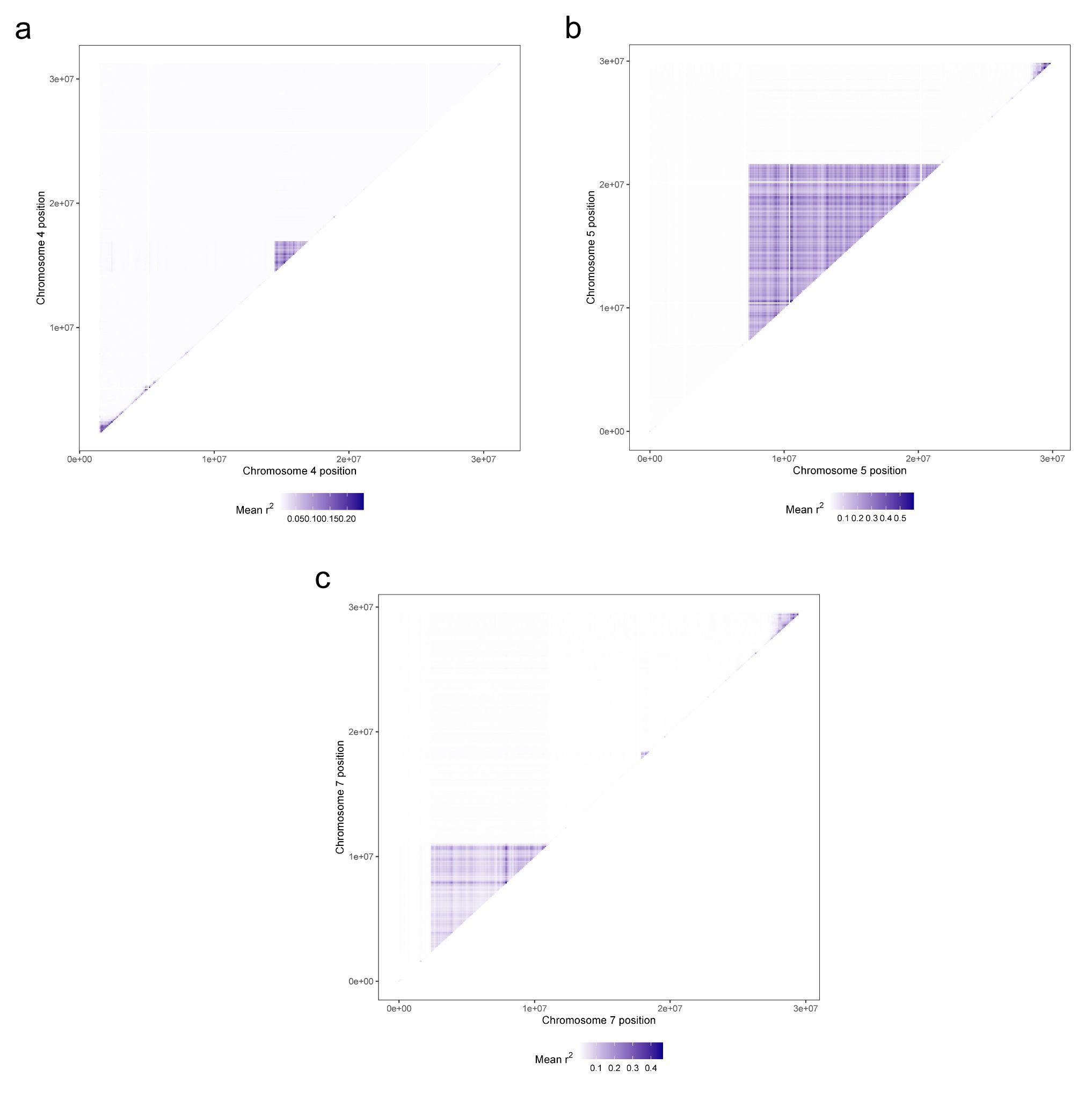

Figure S4. R^2^ heatmaps for plaice chromosomes 4 (a), 5 (b) and 7 (c). Heatmaps for all sole and plaice chromosomes available on dryad.

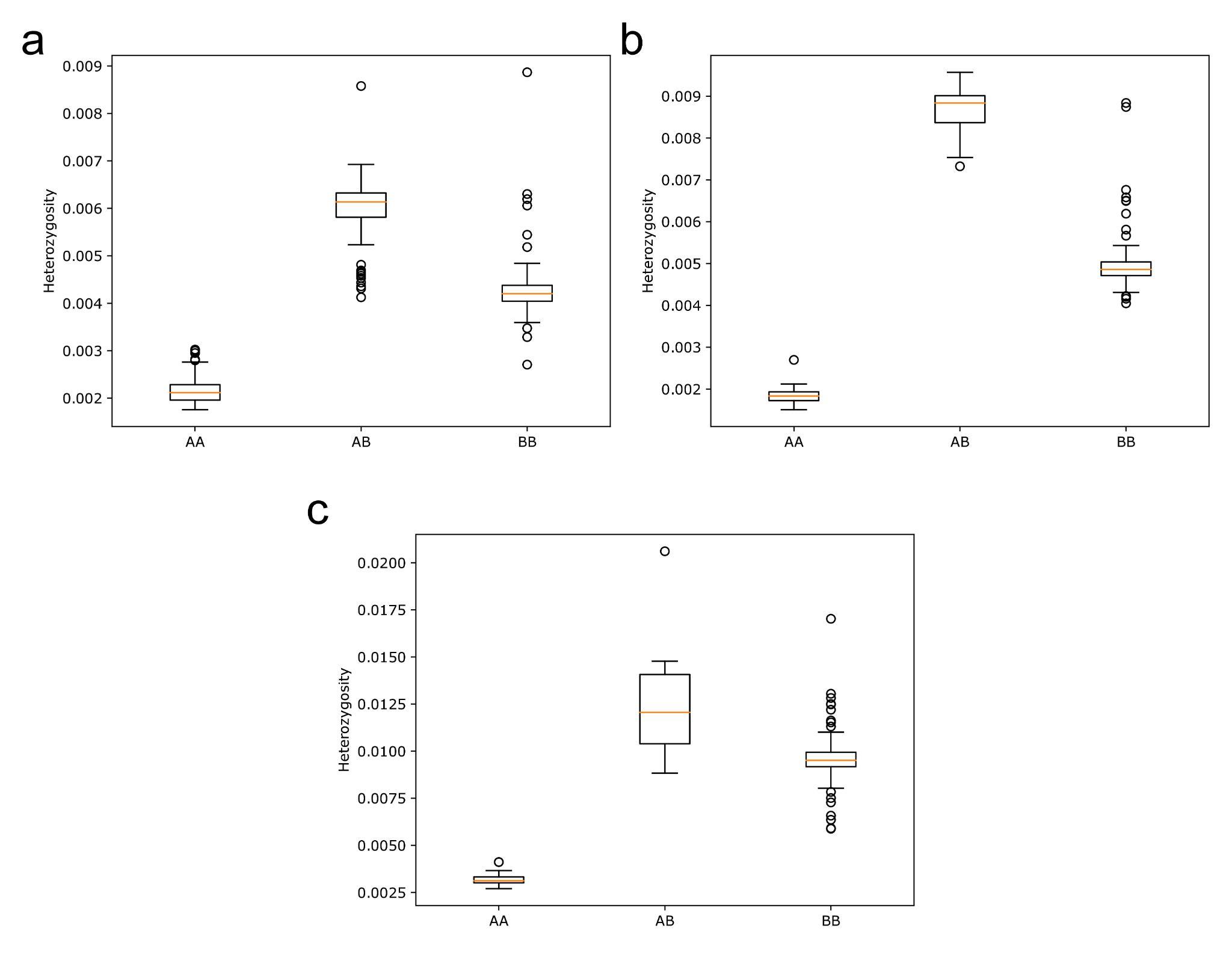

Figure S5. Heterozygosity for individuals with the AA, AB AND BB karyotypes for a) the

chromosome 4 inversion, b) the chromosome 5 inversion; and c) the chromosome 7 inversion.
